# Transcription factor loads tend to decrease the robustness of stable gene transcription networks

**DOI:** 10.1101/037564

**Authors:** Shaoshuai Mou, Domitilla Del Vecchio

## Abstract

Robustness of a system’s behavior to changes in parameter values is a remarkable property of natural systems and especially desirable when designing de novo synthetic gene circuits. Loads on transcription factors resulting from binding to target promoters have been shown to significantly affect the effective time constants of gene transcription networks and to thus alter the overall system’s behavior. Here, we employ models that explicitly account for load effects to investigate how these impact the robustness of a stable gene transcription network to parameter perturbations. By employing a combination of sampling-based methods and analytical tools from control theory, we demonstrate that the presence of loading shrinks the region of parameter space where a gene circuit performs the desired function. A number of multi-module synthetic gene circuits are analyzed to demonstrate this point, including an event detector and a molecular signature classifier. These results indicate that for designing genetic circuits that are robust to parameter uncertainty it is highly desirable to find ways to mitigate the effects of transcription factor loading.

**Author Summary:** Efforts to understand how loads affect gene transcription networks have been underway in the past decade. Here we perform a numerical investigation on three synthetic gene circuits to show that loads tend to decrease the robustness of stable gene transcriptional networks. We complement the numerical findings with analytical derivations that employ the stability radius to compare the robustness of different networks to parameter perturbations near an equilibrium point. Consistent with the numerical finding, the analytical results support that systems with substantial transcription factor loading have smaller stability radius (less robustness) than systems without loading.

## Introduction

Regulation of gene transcription is enabled by the reversible binding reaction of transcription factors (TF) to their target gene promoter sites. It had been theoretically suggested before and experimentally shown later that these binding reactions exert a load on transcription factors, which results in significant effects on both temporal dynamics and steady state [5, 9, 10, 25]. These effects have been called retroactivity to extend the notion of electrical loads to biomolecular systems, making load problems amenable of mathematical study [5]. More recently, the effects of retroactivity have been studied within gene transcription networks (GTN), which result from the regulatory interactions among genes and transcription factors. These studies were performed by employing mathematical models of GTNs that account for the binding of transcription factors to their target promoters [7, 18, 39]. In particular, in [7] the authors have shown that the potential landscape of a toggle switch can be biased by loads. In [39], it was shown that a genetic oscillator can be quenched or robustified depending on what nodes the load is applied to. Finally, in [18] a general ordinary differential equation (ODE) model for gene transcription networks was derived to explicitly account for retroactivity while keeping the same dimension of standard Hill function based models.

Robustness has been studied for a long time in the fields of control, system biology and synthetic biology as a key system property of genetic networks [11–17]. By robustness in this paper is meant the ability to maintain a certain property, such as stability or response time, in the face of parameter perturbations, which may result from genetic mutations [12], changes of interactions among genes [13], or changes in the environment [14]. Robustness enables gene regulatory networks to continue to function despite noisy expression of their constituent genes or even when facing substantial environment variation. From a design point of view, robustness to parameter uncertainty is especially useful as it guarantees that the ideal behavior of a given synthetic circuit is not heavily dependent on the specific parameter values, which are often poorly known.

To study how retroactivity impacts the robustness of gene transcription networks against parameter perturbations, we compare the robustness of two models: the standard Hill function-based model, which does not account for retroactivity [26], and the model developed in [18], which extends the Hill function-based model to include retroactivity. Numerical experiments are then performed on three networks of increasing complexity: a toggle switch, an event detector and a molecular signature classifier. We compare the percentage of success between the system with and without retroactivity when all parameters are sampled in the same intervals. The numerical results indicate that retroactivity leads to more failures in these three systems. To explain this finding more generally, a robustness index called stability radius [29] is introduced to compare local robustness of GTNs with and without retroactivity close to their stable equilibria. Analytical results based on the stability radius also support the finding that retroactivity generally decreases GTNs’ robustness against parameter perturbations.

On the one hand, our finding suggests that natural systems, being inherently robust, may have evolved ways to mitigate retroactivity [19–21]. On the other hand, developing methods to mitigate retroactivity will aid building synthetic biology circuits that are more robust to parameter uncertainty, suggesting that modularity may be instrumental for robustness in addition to being already crucial for bottom-up design [24, 25].

## Models and Problem Formulation

Consider an *n*-node GTN, in which each node has at most two parents. As indicated in Fig.1, each node represents a gene or transcriptional component, whose inputs are the output transcription factors from other nodes. Each directed edge from node *i* to *j* indicated by *i* → *j* in Fig.1 indicates that the output of node *i* regulates the transcription of node *j*. In short, gene transcription networks are composed of “nodes” representing genes and “directed edges” representing regulatory interactions among genes.

Let *x_i_* denote the output transcription factor of node *i* and let *x_i_* denote its concentration. For simplicity of notation we suppose that each node has at most two parents. We will consider and compare the robustness of two models of GTNs. The first one is the standard Hill function-based model [26]. The second model is one that accounts for the binding of TFs to their target operator sites, which is neglected by the standard Hill function-based model [18].

**Figure 1.**
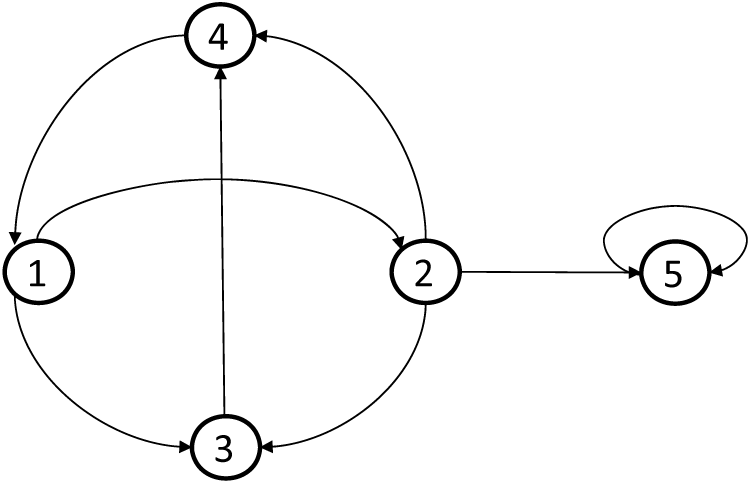
A five-node gene transcription network.

The dynamics of the Hill function-based model can be written as:

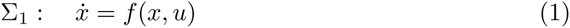

where *x* = [*x_1_ x_2_*… *x_n_*]′ ∈ ℝ^*n*^, *u* = [*u*_1_ *u*_2_… *u_n_*]′ with *u_i_* representing external input to node *i*, and the ith element of *f*(*x, u*) is given by

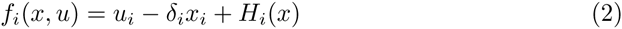

with *δ_i_* denoting the protein decay rate of *x_i_*. Here *H_i_(x)* is the Hill function that models the production rate of *x_i_* as controlled by its two parents *x_p_* and *x_q_* and is given by

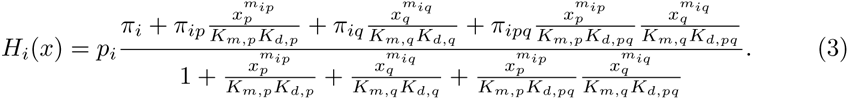

In this expression, *K_m_* is the binding constants for the multimerization; *K_d_* is the binding constants to the promoters; π are specific production rates; *p_i_* is the total concentration of the promoter of node *i*. When all nodes’ *p_i_* are the same, we use *p_T_* to denote this value.

When the effect of the reversible binding between TF and their target promoters is considered, the reaction flux corresponding to this binding reaction appears in the ODE describing the rate of change of the TF’s concentration. This additional flux is what has been called retroactivity [5] and can substantially slow down the temporal response of the TF [9, 25]. Explicitly including this flux in the system’s ODE requires also adding as state variables the concentration of all the complexes that can be formed between promoter sites and TFs, leading to a system with a much higher dimension than that of the Hill function-based model. In [18], leveraging the fact that reversible binding reactions are much faster than the process of gene expression, a reduced model was derived that has the same dimension as the Hill function-based model, yet incorporates the effects of the retroactivity fluxes. According to this model, the dynamics of a gene transcription network modify to:

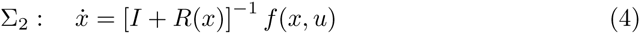

where *R(x)* ∈ ℝ*^n × n^*, called the *retroactivity matrix*, is given as

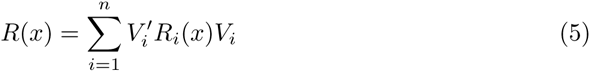

where *V_i_* ∈ ℝ^*n_i_* × *n*^ with *n_i_* the number of node *i*’s parents. The *jk*th element of *V_i_* is 1 if the *j*th parent of *i* is *k*, and is 0, otherwise. We call *R_i_(x)* the retroactivity of node *i* and will be discussed in more details in the following.

The Hill function *H_i_(x)* and the retroactivity of node *i R_i_(x)* depend on the number of node *i*’s parents and the bindings with its parents [18]. If node *i* has no parent, one has 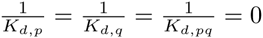 in (3) and *R_i_(x)* = 0; if node *i* has a single parent node *x_p_*, we let 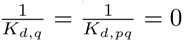 in (3), and

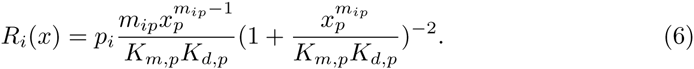

When node *i* has two parents x_*p*_ and x*_q_*, the values of *H_i_(x)* and *R_i_(x)* depend on the binding type, which is typically one of the following:

- *Competitive binding:* x*_p_* and x*_q_* bind exclusively to the promoters of their common child. In this case, one has 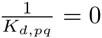 in (3) and

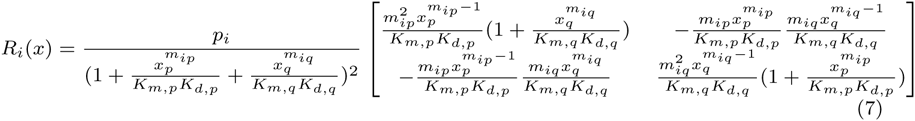
- *Independent binding:* x_*p*_ and x_*q*_ do not affect each other in their bindings to a common child. That is, even if a node’s promoter is bound with one parent, it is still available to be bound with its other parents. In this case, *H_i_(x)* is as defined in (3) and

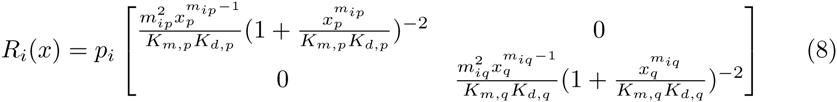
- *Cooperative binding:* x*_p_* must be bound to its child’s promoters before x*_q_* can bind. In this case, one has 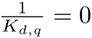 in (3) and

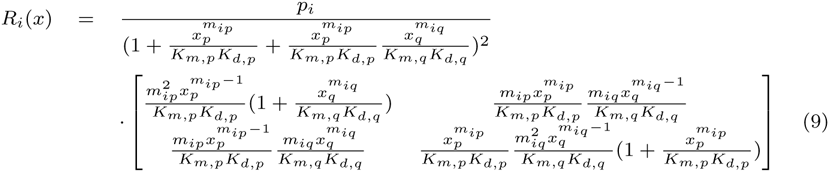 For details on these derivations, the reader is referred to [18].

## Numerical Experiments

In this section numerical experiments are performed to show the impact of retroactivity on the robustness of a toggle switch, an event detector and a molecular signature classifier to parameter variations. This is performed by comparing the parameter spaces where the desired behavior is obtained for model Σ_1_ (without retroactivity) and model Σ_2_ (with retroactivity). For simplicity we only consider the case where all promoters have the same total concentration *p_T_*, modeling the case in which the systems’ parts are inserted all in the same plasmid with concentration *p_T_*.

### Toggle Switch

We first consider the toggle switch, which is a simple module exhibiting bistable behavior, originally constructed in [1] and then used as a switch in many other more sophisticated multi-module systems [2, 3]. As shown in Fig. 2, the toggle switch is composed of two TF x_1_ and x_2_, which negatively regulate each other’s transcription. Here, *i* ⊣ *j* indicated that node *i* is repressing node *j*. Under suitable conditions (see [4], for example) the toggle switch has two stable steady states, at each of which, one of the TF appears in a high copy number, while the other one is suppressed.

**Figure 2.**
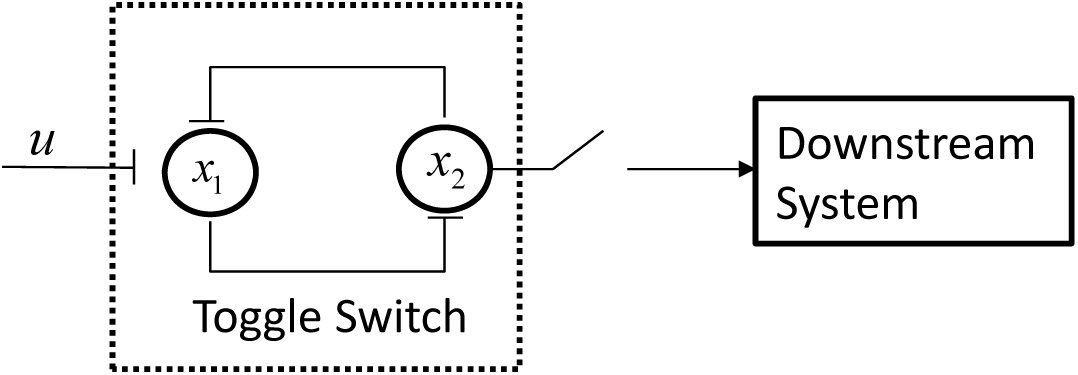
The Toggle Switch.

With input *u* regulating x_1_, the dynamics of the toggle switch without retroactivity Σ_1_ are given by

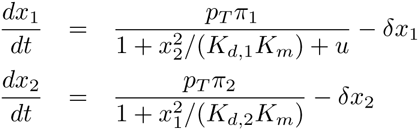

in which we have assumed for simplicity that all the transcription factors have the same half lives. The dynamical model of the toggle switch with retroactivity Σ_2_ is given by

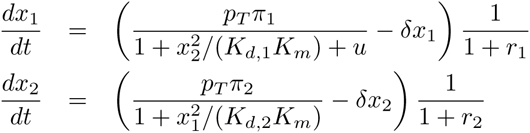

where, from expression (6), we have that

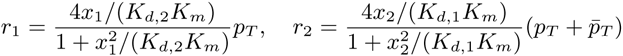

Here 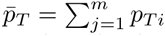 and M*p_Ti_* denotes the total concentration of x_2_’s jth child promoter contained in the downstream system. When the toggle switch is not connected to any downstream system, 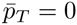.

For a given input profile u switching from a low value to a high value, the toggle switch is said to be functional if the output *x*_2_ switches from low to high and keeps this high value even when the input u changes back to its low value. If the toggle switch output does not switch to its high value and latches to it, the toggle is said to be non-functional. As a demonstration of the effect of retroactivity on the toggle switch dynamics, we illustrate numerical simulations in Fig. 3. The system model without retroactivity Σ_1_ is functional. When the toggle switch is not connected to any downstream system, that is, 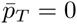, the system model Σ_2_ also functions (as indicated in Fig. 3(a)) but the switching time increases, that is, the system becomes slower. This is in accordance to what demonstrated in previous studies [8, 18]. When the toggle switch is connected to a downstream system, it fails to function as no switch is observed (Fig. 3(b)). If in this case, one increases the decay rate, the switching is restored even if the final value is lower (Fig. 3(c)). This is in accordance with the fact that if the temporal response of a TF is faster (as obtained, for example, by increased turnover rates), retroactivity has a decreased effect [25].

**Figure 3.**
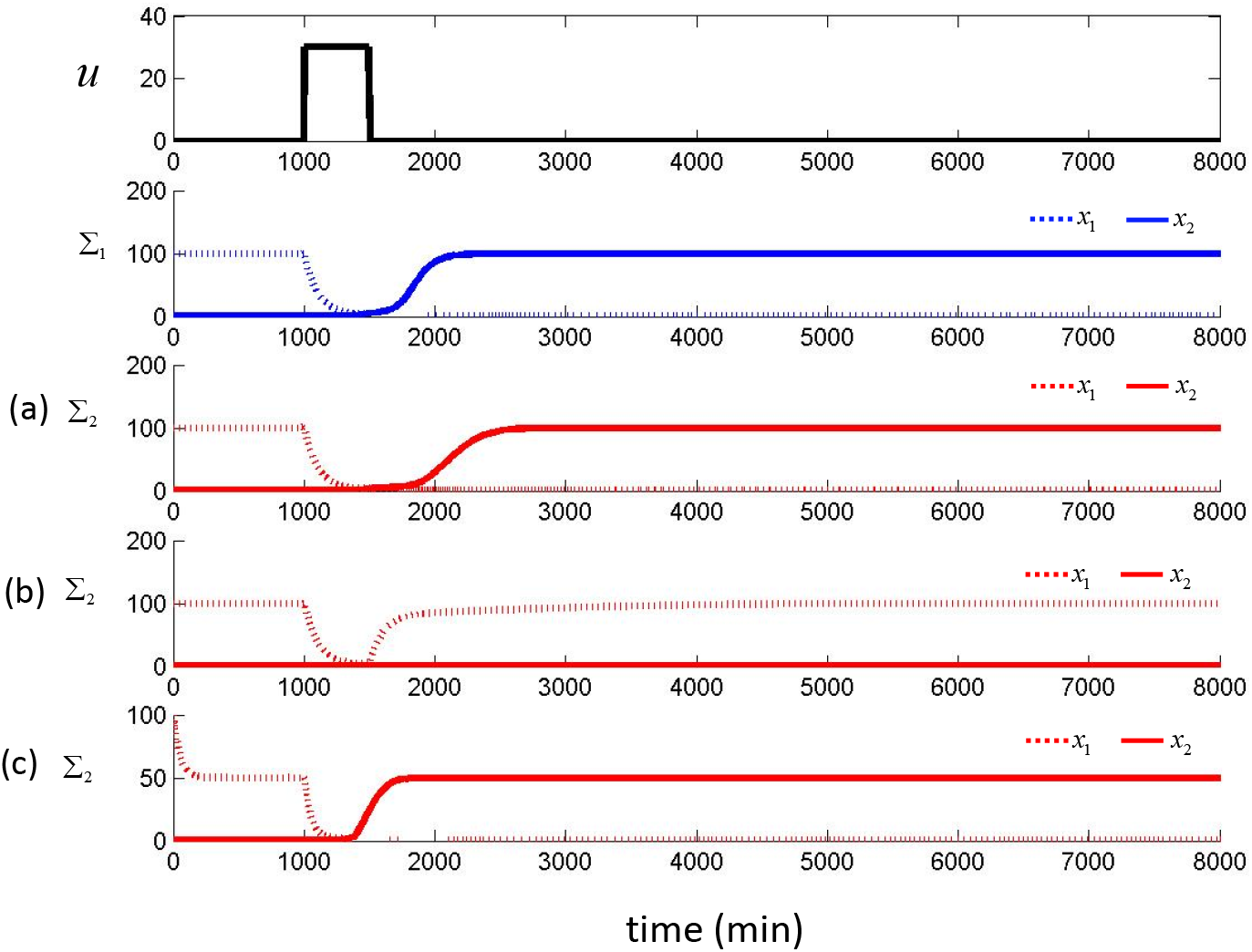
Effect of retroactivity on the toggle switch. Comparison between Σ_1_ and Σ_2_ for fixed parameters. Here, we have set δ = 0.01 min^−1^, *K*_*d*, 1_ = *K*_*d*, 2_ = *K_m_* = 1 nM, *p_T_* = 1 nM, π_1_ = π_2_ = 1 min^−1^. The system without retroactivity Σ_1_ is functional. (a) Model with retroactivity Σ_2_ without downstream system 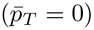. (b) Model with retroactivity Σ_2_ with downstream system 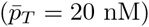. (c) Model with retroactivity Σ_2_ with downstream system 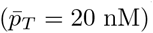 but with increased decay rate δ = 0.02.

### Robustness to parameter variations

To determine how retroactivity affects the robustness of the toggle switch to parameter variations, we compare the fraction of parameter space for Σ_1_ and Σ_2_ that leads to a functional toggle switch. A larger fraction of the parameter space leading to a functional system indicates larger robustness to parameter variations.

To this end, we treat each parameter as an independent random variable uniformly distributed in a certain interval. Parameters for concentrations are in nM and time is in minutes. In this section, we choose *δ* ∈ [0.01,0.02], which means that the half life of proteins is in [30, 60] minutes. Choose *p_T_* ∈ [1, 100] to include both low and relatively high plasmid copy number. By considering that in practice the number of copies of each protein per cell is expected to be less than 20000, we choose π_i_ such that 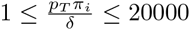. Then one has π_i_ ∈ [0.02, 2]. Suppose the disassociation constants *K_d_* and *K_m_* are in [1, 50] (see, for example [18]). For each fixed 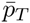, we employ the Latin Hypercube Sampling (LHS) method to take a number of N samples of all the other parameters from their corresponding intervals and run simulations of Σ_1_ and Σ_2_ for each sample. LHS is a type of stratified Monte Carlo sampling method, which is highly efficient. In fact, in practice it is sufficient that *N* is larger than 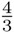 times the number of random variables to provide a statistically sufficient covering of the entire parameter space [27]. Here, we choose *N* = 2000, which is much larger than what found to be sufficient for LHS. For the same input profile *u(t)* as given in Fig. 3, if a switch in the output of x_2_ is observed, we call it a success; otherwise, we call it a failure. We count the number of successes of Σ_1_ and Σ_2_ and summarize the results in the percentages shown in Fig. 4.

**Figure 4.**
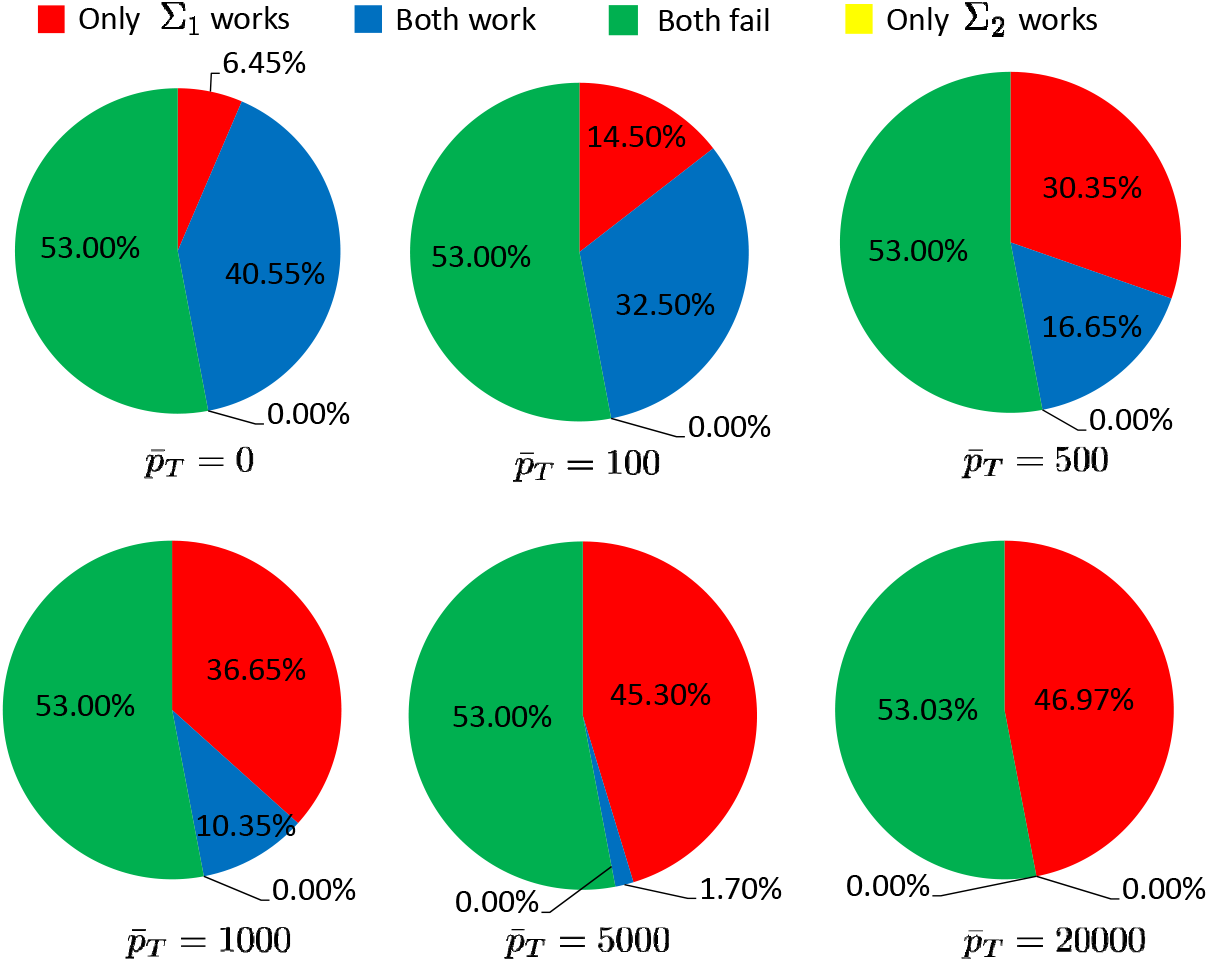
Robustness of toggle switch to parameter perturbations. Percentage of the parameter space that leads to success of Σ_1_ (without retroactivity) and Σ_2_ (with retroactivity) shown in Red plus Blue and Blue, respectively. Here, 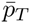 corresponds to the number of promoter sites in the downstream system that x_2_a regulates.

Fig. 4 shows that the system without retroactivity Σ_1_ always has a larger fraction of the parameter space leading to success when compared to the system with retroactivity Σ_2_. In particular, as the number of promoter sites 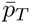 that x_2_ regulates in the downstream system increases, the fraction of the parameter space where the system with retroactivity functions shrinks to the point of never functioning when 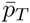 is extremely high. Note that the percentage of cases where the system with retroactivity functions and the one without retroactivity does not is zero.

### Event Detector

In this section, we perform simulations on an event detector circuit and illustrate how retroactivity affects the robustness of such a multi-module system against parameter perturbations. The event detector (ED) consists of six nodes and is shown in Fig. 5, in which *i* → *j* and *i* ⊣ *j* represent that i is an activator or a repressor of *j*, respectively. The event detector detects a decrease in the input by switching the value of the output node to a low value and by keeping it even after the input has acquired back the original high value (by virtue of the toggle switch). In particular, when the input *u* switches to a low value, the cascade consisting of nodes x_1_, x_2_, x_3_ propagates the signal to remove repression on the inverter x_4_, eventually resulting in a switch in the state of the toggle module, which leads to a switch of the output node x_7_ to a low value.

**Figure 5.**
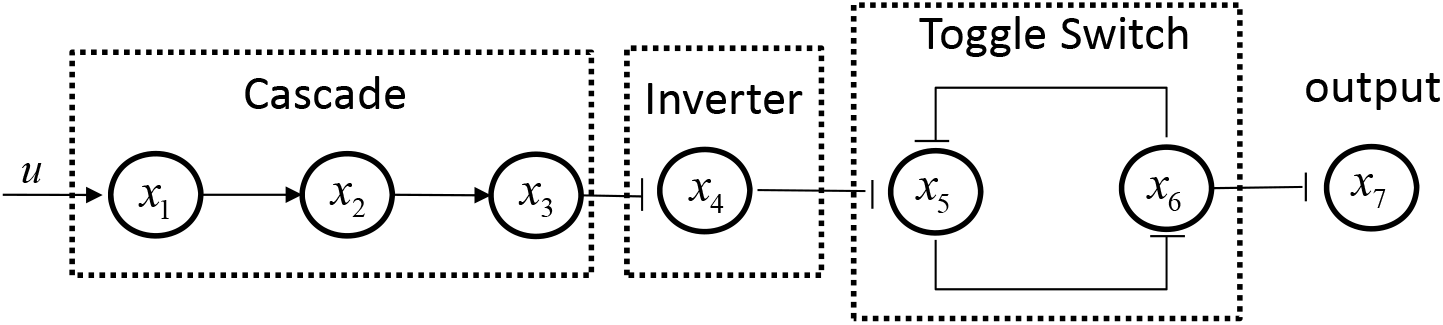
The Event Detector.

The dynamics of the event detector without retroactivity are given by system model Σ_1_: 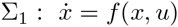 where *x* = [*x*_1_ *x*_2_ *x*_3_ *x*_4_ *x*_5_ *x*_6_ *x*_7_]′ and the *i*th element of *f (x, u)* is given as follows:

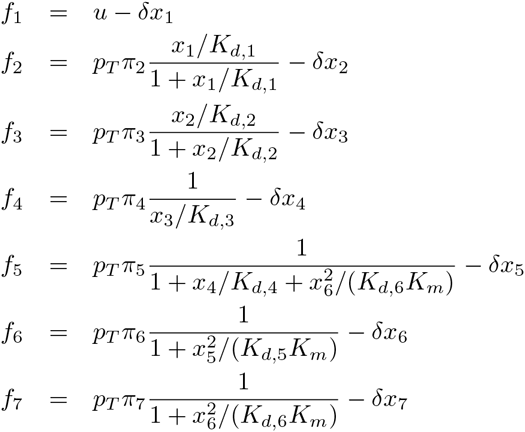

By considering retroactivity, one has system model Σ_2_:

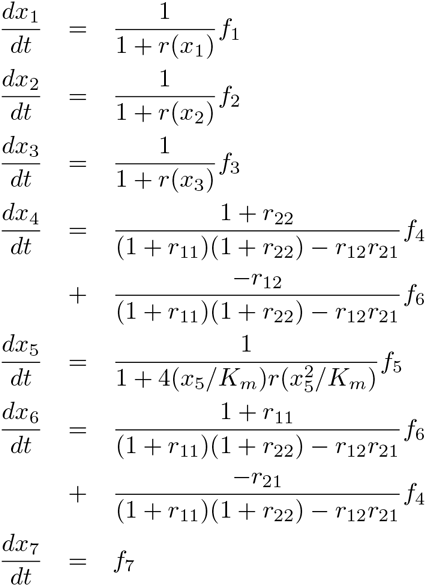

where *r(X)*, *r*_11_, *r*_12_, *r*_21_, *r*_22_ are the retroactivity expressions given by

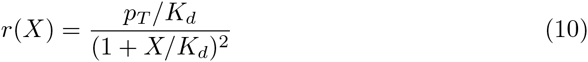

and

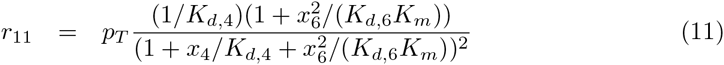

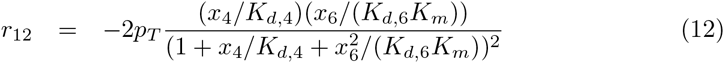

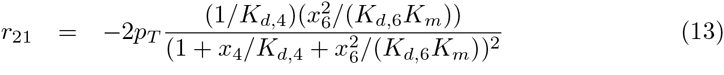

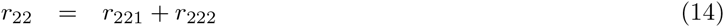

with

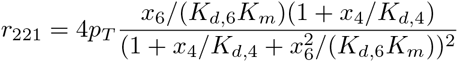

and

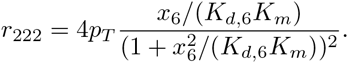

Here, *r*_11_, *r*_12_, *r*_21_, *r*_221_ follow from (7) since x_4_ and x_6_ bind competitively to their common child x_5_, and *r*_222_ follows from (6). Note that when all the above retroactivity expressions are 0, Σ_2_ is exactly the same as Σ_1_.

For a given input profile u that switches from a high value to a low value, the ED is said to be functional if the output *x*_2_ switches from high to low and keeps this low value even when the input *u* changes back to its high value. If the ED’s output does not switch to its low value and latches to it, the system is said to be non-functional. As a demonstration of the effect of retroactivity on the ED’s dynamics, we illustrate numerical simulations in Fig.6.

The output of the ED without retroactivity Σ_1_ indicated by blue plots in Fig. 6 successfully detects the event while the ED with retroactivity Σ_2_ fails. As we progressively increase δ, both Σ_1_ and Σ_2_ work properly. This reaffirms the fact that retroactivity has less of an impact on TF with faster dynamics as described in the case of the toggle switch. In fact, it is well known that retroactivity leads to delays in the temporal response of transcription factors [9, 25], which are accumulated through the stages of the cascade as illustrated in Fig. 7, ultimately leading to the ED’s failure. A faster TF turn over rate mitigates the effects of load-induced delays [22, 25].

### Robustness to parameter variations

To determine how retroactivity affects the robustness of the ED to parameter variations, we compare the fraction of parameter space for Σ_1_ and Σ_2_ that leads to a functional ED. A larger fraction of the parameter space leading to a functional system indicates larger robustness to parameter variations.

To this end, we randomly change all parameters in Σ_1_ and Σ_2_ and check whether the ED still functions. We employ as before LHS to select 2000 samples of parameters from intervals δ ∈ [0.01, 0.02], π_*i*_ ∈ [0.02, 2], *p_T_* ∈ [1, 100], *K_d_* and *K_m_* in [1, 50]. If the output of the event detector is able to detect the change of the input and maintain it at the low value, we count it as a success. Results are summarized in Fig. 8. The percentage of parameter space where Σ_1_ successfully functions (red+blue) is 79.75% while that where Σ_2_ functions (blue) is only 42.20%. That is, the system model with retroactivity successfully functions in about only half of the parameter space where the system model without retroactivity functions.

**Figure 6.**
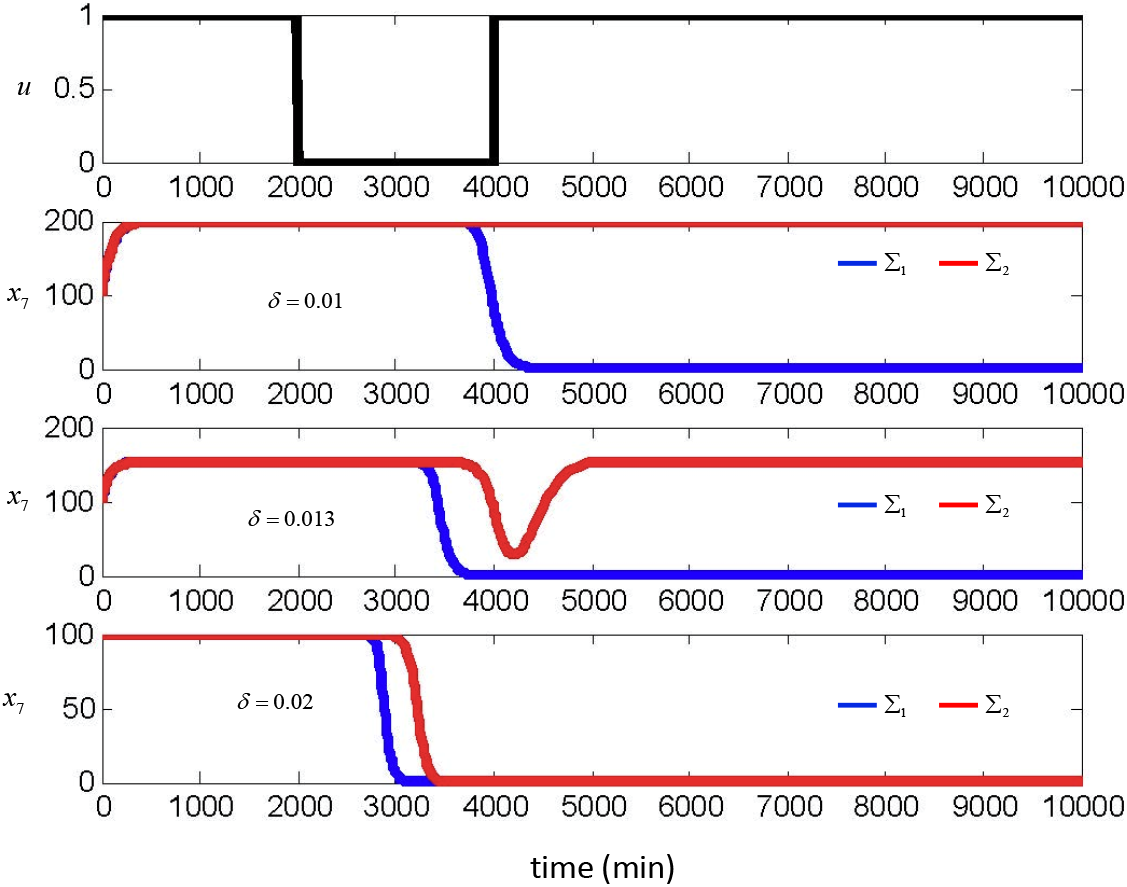
Effect of retroactivity on the event detector. The output of Σ_1_ (without retroactivity) and Σ_2_ (with retroactivity). Parameter values are set to *p_T_* = 1nM, *K_m_* = 10nM, *K_d, i_* = 1nM, π_i_ = 2min^−1^. The value of *δ* is as indicated in the plots.

To further determine the relationship between the circuit copy number *p_T_*, which determines the load applied by target promoters on their transcription factors, and the failure due to retroactivity, we then fixed *p_T_* at the different values 1, 5,10, 20, 50,100 and randomly changed all the other parameters in their respective intervals. Results are summarized in Fig. 9. When the total concentration *p_T_* is low, the failure due to retroactivity is only 0.1%, which implies that Σ_1_ and Σ_2_ behave similarly and retroactivity does not have a dramatic impact. By contrast, when *p_T_* is increased to 100, the failure due to retroactivity grows to 64.60%. That is, the existence of retroactivity causes 64.60% of the parameter space of the event detector to lead to a non-functional system. All together, these results indicate that retroactivity dramatically decreases the robustness of the ED to parameter variations and that an suggest that an ED built on very low plasmid copy number (*p_T_*) will be more robust to parameter variations. Of course, tradeoffs with noise may become important as the molecule count decreases.

### Classifier

In this section, we consider a molecular signature classifier circuit that is composed of five modules as shown in Fig. 10. These modules are three sensors, an AND gate whose design is based on [6], and the toggle switch [1]. The output of the classifier should be switched OFF shortly after all the three inputs *u_1_*, *u_2_*, *u_3_* have become high at the same time. Here, the inputs *u*_1_, *u*_2_, *u*_3_ represent the concentrations of three different signaling molecules and the situation of interest is when they are all high simultaneously. As soon as the inputs become all high simultaneously, the concentrations of TF x_6_, x_1_, x_5_, and thus of x_2_ become high. Since TF x_3_ and x_4_ are activated by a cooperative interaction of their parent nodes, the concentration of x_4_ becomes high when and only when all the three inputs *u*_1_, *u*_2_, *u*_3_ are high at the same time. TF x_4_, in turn represses x_7_. Therefore, the toggle switch TF x_8_ switches to its high value and turns OFF the output.

The dynamics of the above classifier without retroactivity are 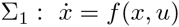, where *x* = [*x*_1_ *x*_2_ *x*_3_ *x*_4_ *x*_5_ *x*_6_ *x*_7_ *x*_8_ *x*_9_]′ and the *i*th element of *f(x, u)* is given as follows:

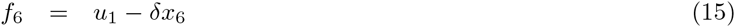

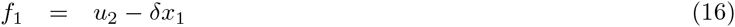

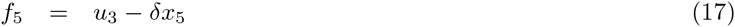

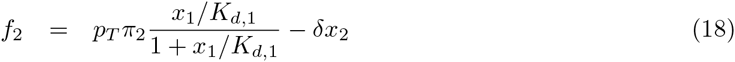

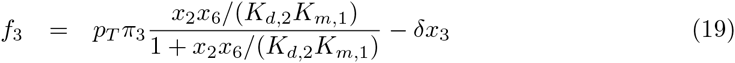

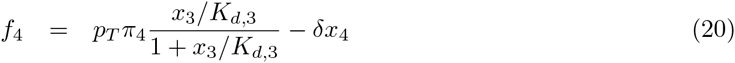

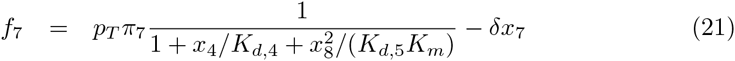

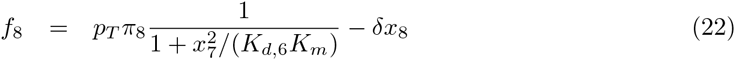

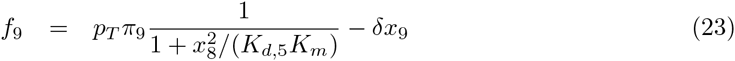

**Figure 7.**
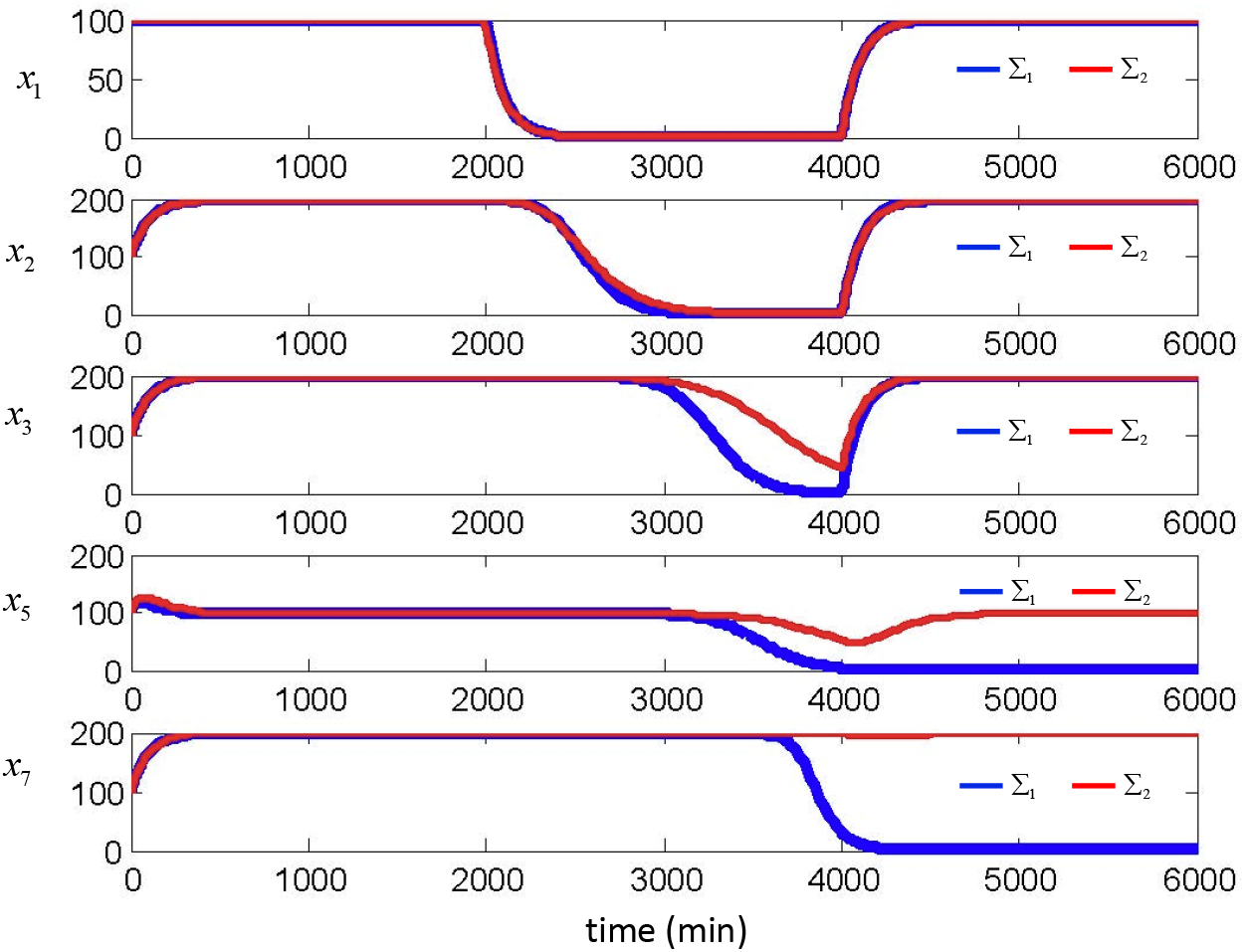
Propagation of load-induced delays in the event detector. Comparison of Σ_2_ (with retroactivity) and Σ_1_ (without retroactivity).

By considering retroactivity one has a different model denoted by Σ_2_ described by the following ODEs:

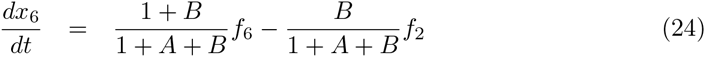

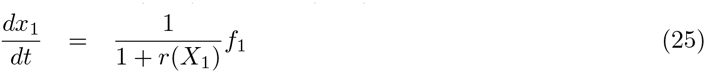

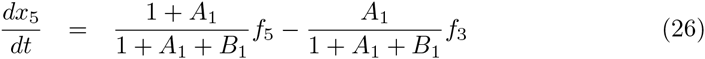

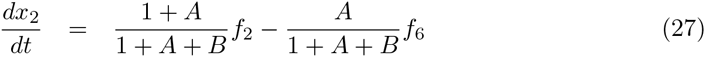

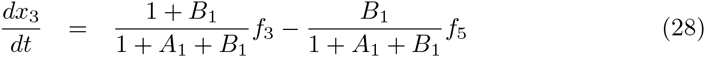

**Figure 8.**
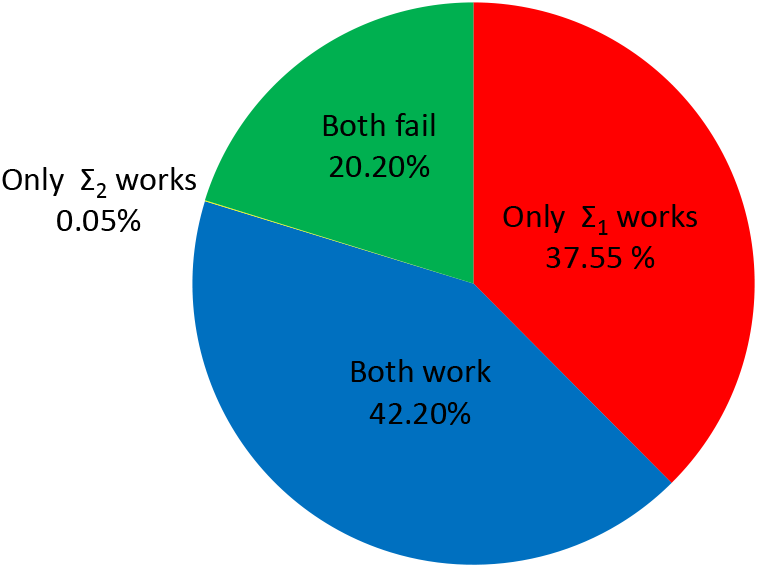
Robustness of the event detector to parameter perturbations. Percentage of the parameter space that leads to success of Σ_1_ (without retroactivity) and Σ_2_ (with retroactivity) shown in Red plus Blue and Blue, respectively.

**Figure 9.**
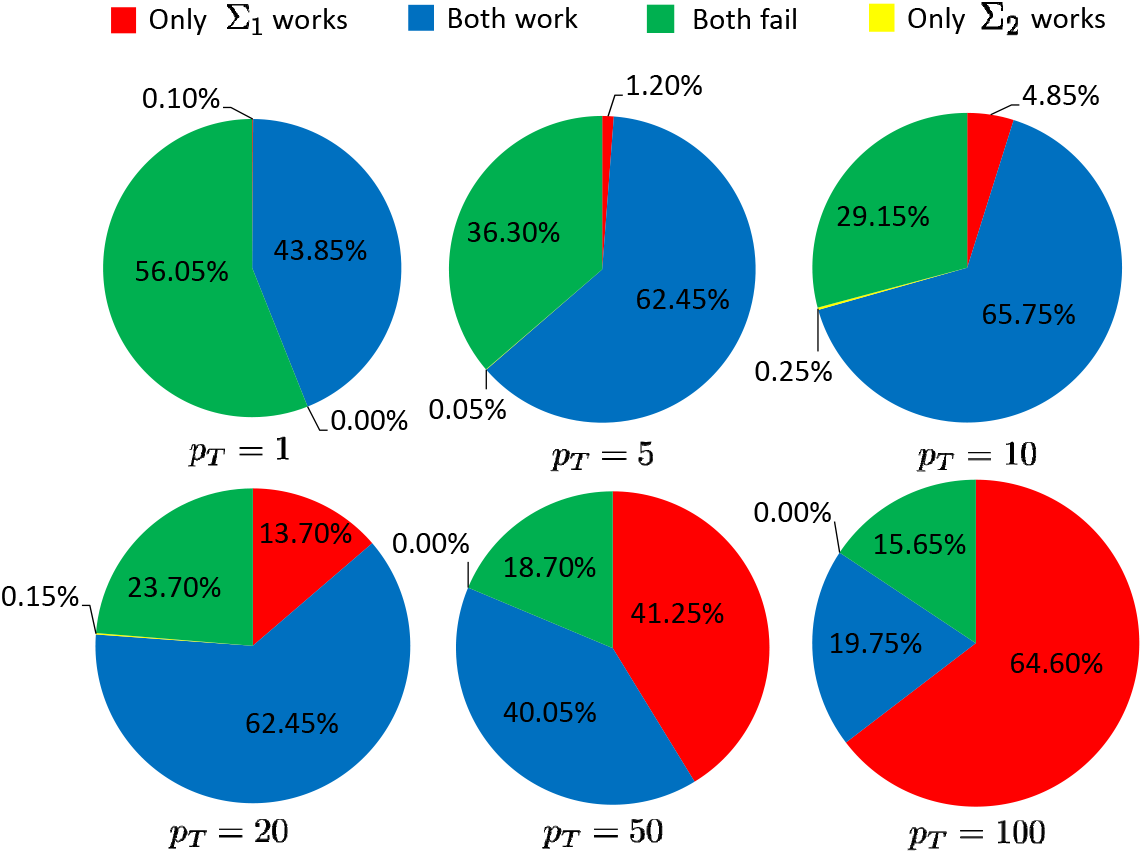
Percentage of success of Σ_1_ (without retroactivity) and Σ_2_ (with retroactivity) for different values of *p_T_*.

**Figure 10.**
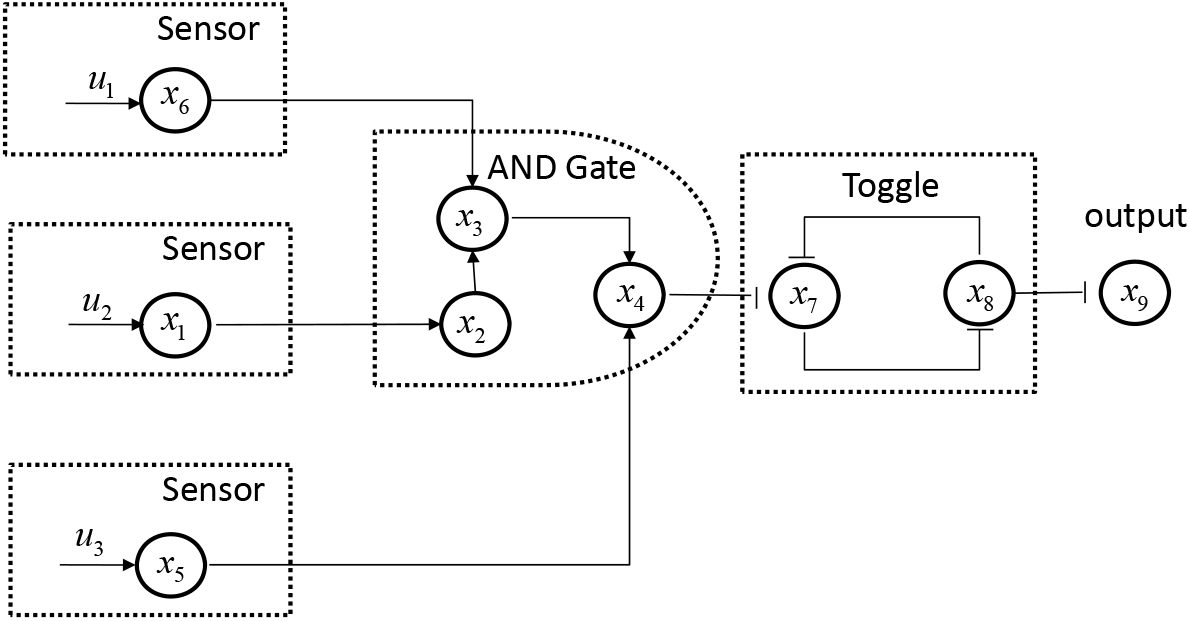
Classifier

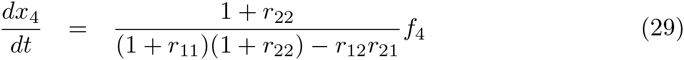

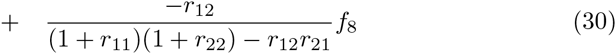

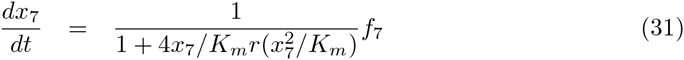

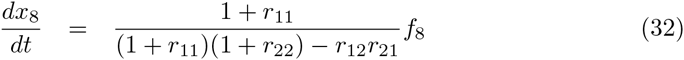

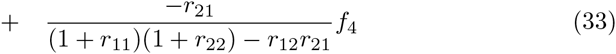

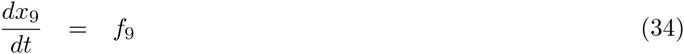

Here, *r(X)*, *r*_11_, *r*_12_, *r*_21_, *r*_22_ are the same expressions as (10)-(14); *A, B, A*_1_ and *B*_1_ are due to cooperative bindings of x_6_ and x_2_ to x_3_, and x_3_ and x_5_ to x_4_, respectively, for which one has by (9)

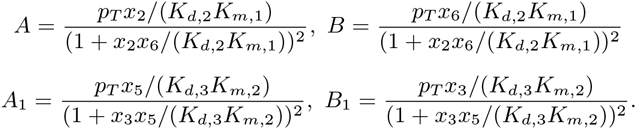

For a given triple of input profiles *u*_1_, *u*_2_, *u*_3_, the classifier is said to be functional if the output *x_9_* switches from high to low when and only when the inputs *u_i_* become all high simultaneously. If the classifier’s output does not switch to its low value and latches to it, the system is said to be non-functional. As a demonstration of the effect of retroactivity on the classifier’s dynamics, we illustrate numerical simulations in Fig. 11. For the given input profile *u*_1_, *u*_2_, *u*_3_, the output of the classifier without retroactivity Σ_1_ indicated by the blue plots successfully switches from high to low as soon as all the inputs become high simultaneously. By contrast, system Σ_2_ fails since its output becomes low even when the input *u*_2_ is still low. The reason for which Σ_2_ fails is the accumulation of time delays in the temporal response of transcription factors caused by retroactivity. This phenomenon also agrees with the observation of delay’s effects in Fig. 7. Note from the red plots of Fig. 11 that the time delay experienced by the pulse resulting from *u*_2_ propagates to *x*_3_, so that the AND of *x*_3_ and *u*_4_ results into a high x_4_ that ultimately switches the output of the toggle switch OFF, leading to failure of the classifier. Once we change δ to 0.02, both Σ_1_ and Σ_2_ function as this delay does not accumulate as much. This is consistent with the previous observations that retroactivity has less of an impact if the TF have faster turnover rates.

**Figure 11.**
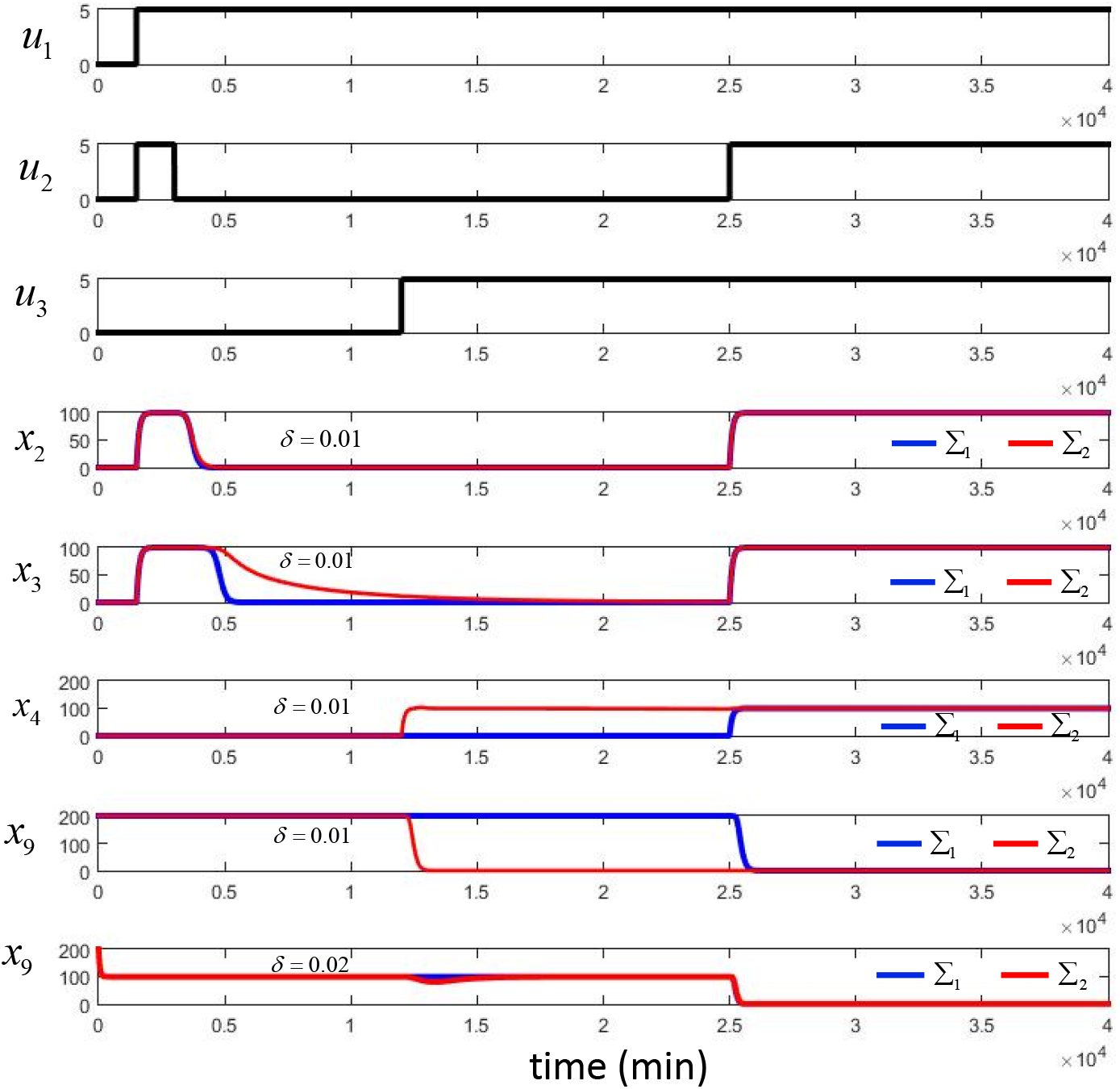
Comparison between the Classifier Σ_1_ without retroactivity (indicated in blue) and the one with retroactivity Σ_2_ (indicated in red). Parameter values are given by *p_T_* = 1 *K*_*d*, 1_ = *K*_*d*, 2_ = *K*_*d*, 3_ = *K*_*d*, 5_ = *K*_*d*, 6_ = 1, *K*_*d*_, 4 = 10, *K_m_* = *K*_*m*, 1_ = *K*_*m*, 2_ = 10, π_1_ = Π_2_ = Π_3_ = Π_4_ = Π_5_ = 1 and π_6_ = 2.

### Robustness to parameter variations

To determine how retroactivity affects the robustness of the classifier to parameter variations, we compare the fraction of parameter space for Σ_1_ and Σ_2_ that leads to a functional classifier. A larger fraction of the parameter space leading to a functional system indicates larger robustness to parameter variations.

To this end, we employ the LHS method as before to obtain 2000 samples from the parameter space. For each sample of parameters, we employ the inputs *u*_1_, *u*_2_, *u*_3_ as shown in Fig. 11. If the output of the classifier at x_9_ is low if and only if all *u*_1_, *u*_2_, *u*_3_ are high, we count it as a success; otherwise, it is counted as a failure. Simulation results including the failure due to retroactivity (marked as red) are shown in Fig. 12, which suggests that retroactivity leads to malfunction in 47.40% of the parameter space.

To further determine the relationship between *p_T_* and the loss of function of the classifier, we fixed *p_T_* = 1, 5,10, 20, 50,100 and randomly changed all the other parameters. Simulations are summarized in Fig. 13. When the total concentration *p_T_* is low, for example *p_T_* = 1, the failure due to retroactivity is 0.05%, which implies that Σ_1_ and Σ_2_ behave similarly and retroactivity does not have dramatic impact ono the classifier. When p_T_ is increased to be 100, one observes a large value of the failure due to retroactivity, which is 70.05%.

**Figure 12.**
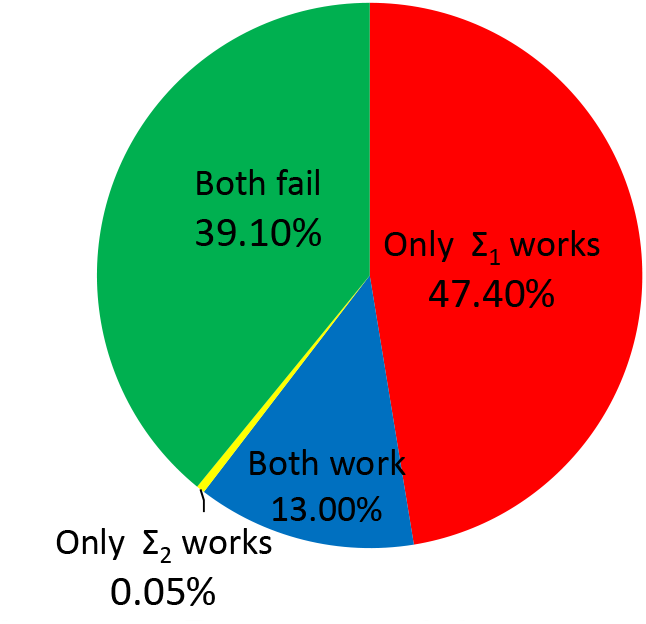
Classifier robustness. Percentage of the parameter space that leads to success of Σ_1_ (without retroactivity) and Σ_2_ (with retroactivity) shown in Red plus Blue and Blue, respectively.

**Figure 13.**
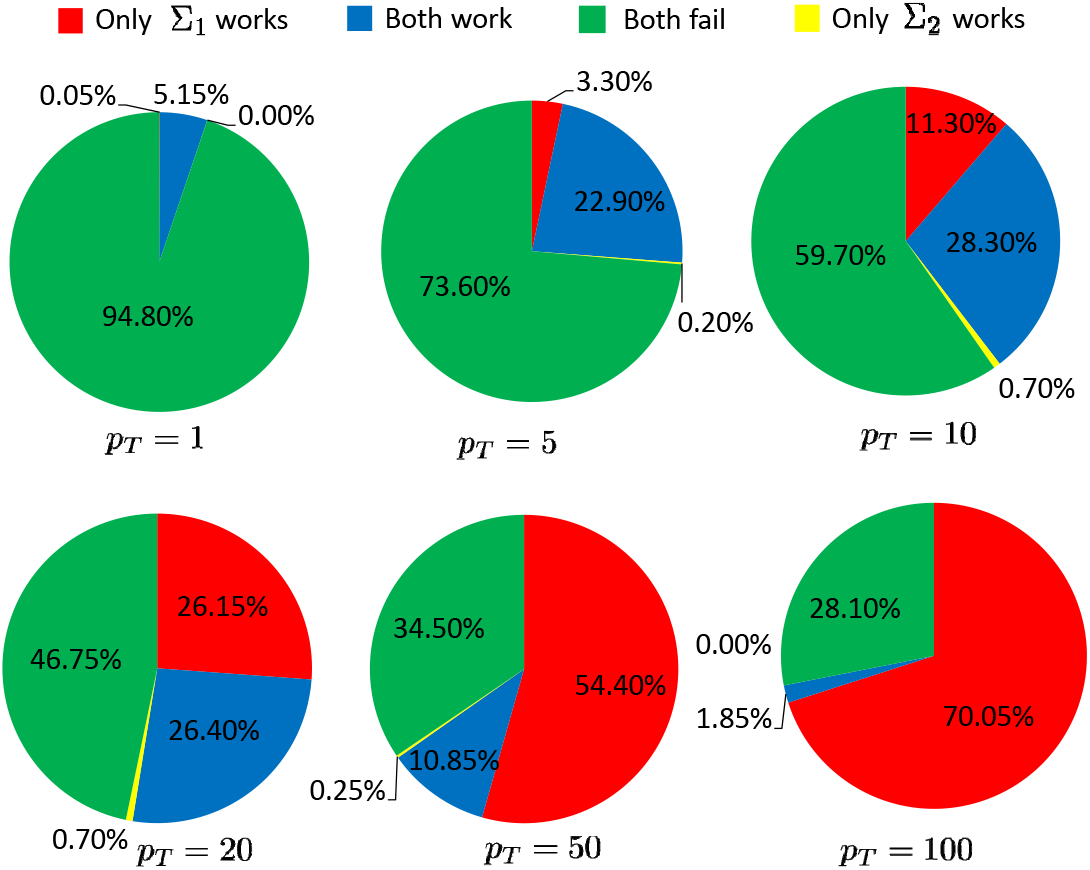
Percentage of success of the classifier Σ_1_ (without retroactivity) and Σ_2_ (with retroactivity) for different values of *p_T_*.

### Summary of Findings from Numerical Simulations

Simulations on the toggle switch, the event detector and the classifier suggest the following. When we sample the parameter space for Σ_1_ and Σ_2_, system with retroactivity Σ_2_ encounters substantially more failures than the system without retroactivity Σ_1_. This indicates that retroactivity shrinks the region of parameter space where these stable gene circuits perform the desired function and thus decreases their robustness. In accordance to what previously found, retroactivity’s impact on the system’s robustness is more dramatic on a slower system than on a faster one. When a system is fast enough, the impact due to retroactivity becomes negligible.

## Analytical Measure of Robustness

In this section, we analytically compare the robustness of the system with retroactivity to that of the system without retroactivity and thus confirm more generally that retroactivity tends to decrease the robustness of stable gene transcription networks against parameter perturbations. To this end, we analyze the behavior of the systems (with and without retroactivity) close to the common stable equilibrium *x*\*, where *x*\* is such that *f(x*, u)* = 0 for a fixed *u*. The objective is to compare the robustness of the equilibrium’s stability to parameter perturbations. We thus consider the linearization of Σ_1_ and Σ_2_ about *x*\*, which leads to the two following linear systems:

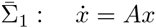

and

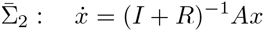

where

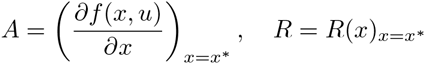

are constant matrices. For the system’s robustness with respect to parameter perturbations, we restrict ourselves to additive perturbations, which, compared to relative perturbations that inherently have a multiplicative structure, are the most general [28]. To mathematically compare the robustness of 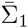 and 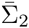 against additive parameter perturbations, we employ the concept of stability radius, which has a long history in robust control theory [29, 30]. The stability radius measures a system’s ability to maintain certain stability conditions of the equilibrium point under additive perturbations to the elements of the system’s matrix. If system 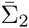 has smaller stability radius than system 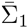, it follows that the worst case parameter perturbation in 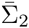 pushes the slowest eigenvalue closer to the imaginary axis than the worst case parameter perturbation does in 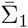. As a consequence, we should expect much slower convergence to the equilibrium in 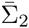 as compared to 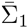 in the worst case. In the sequel, we will say that 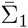 is more robust than 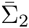 if the stability radius of the former is greater than that of the latter.

In particular, let Λ(*M*) denote the spectrum of a square matrix *M* ∈ 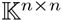^*n × n*^, where 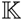 = ℂ or ℝ. Let ℂ^−^ denote the open left-half complex plane and let ℂ^+^ denote the closed right-half complex plane. Define the *stability radius* of *M* as

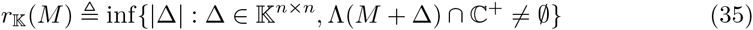

where | · | denotes the 2-norm. Then 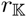(*M*) is the 2-norm of the smallest perturbation forcing *M* + Δ to be unstable. The stability radius defined in (36) is a natural measure of a system’s ability to maintain stability of an equilibrium point under perturbations to elements of the system matrix. A system with larger stability radius is able to maintain its stability under larger perturbations to the system’s matrix in the 2-norm sense.

The computation of *r*_ℝ_(*M*) is in general a challenging problem [32]. To avoid complex computations, we determine lower and upper bounds of *r*_ℝ_(*M*) for both the system with retroactivity 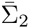 and the system without retroactivity 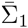, which can be more easily computed, and then compare the bounds. The derivations of the bounds (see the Appendix) can be easily performed when the retroactivity matrix *R(x)* is diagonal, corresponding to the case in which transcription factors bind to the corresponding target promoters independent of each other or almost independent (i.e., the off-diagonal entries of *R(x)* are sufficiently small compared to the diagonal entries). In such a case, let 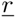 and 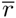 represent the smallest and largest diagonal entries of *R*, respectively. Then, we can analytically prove (see Appendix) that 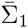 is more robust than 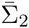 when any of the following cases holds:

i. 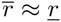, that is, the diagonal entries are all close to each other;
ii. 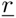 is large enough, that is, retroactivity is high;
iii. 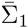 has some low-pass filtering properties in the sense that the *H*_∞_ norm of the matrix *A* is achieved at *ω* = 0.

Therefore, if the loads resulting at all nodes of the network are balanced (i.e., they are close to each other), if the loads at all nodes are very large, or if the system response to higher frequencies stimulations is lower compared to that at low frequencies (low-pass filtering behavior), which is often the case in biomolecular networks ([34-36]), then 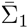 is more robust than 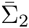.

## Discussion

In this paper, we have analyzed how the robustness of a stable gene transcription network is affected by the loading applied on transcription factors by the promoter sites to which the factors bind. Here, robustness is intended as the ability of a system to keep a desired behavior in the face of parameter perturbations. Specifically, the behavior whose robustness is investigated is the stability of equilibria, which is especially important in systems with memory, including switches, event detectors, and molecular signature classifiers. These enable a cell to make a decision based on changes in the molecular profile of the environment and have been extensively investigated for synthetic biology applications [40].

Our computational study performed by sampling the biologically relevant parameter space indicates that the parameter region where the desired behavior is observed is more than 46% smaller in the system model that includes retroactivity compared to one that does not include it. Since the impact of retroactivity is controlled by the dissociation constant of TF to their promoter sites and by the promoter sites number (DNA copy number), we also studied how the parameter region corresponding to the desired behavior is affected by increasing the DNA copy number. For low copy numbers, the two models (with and without retroactivity) have similar parameter regions leading to the desired behavior (less than 10% difference). However, for medium and high copy numbers this parameter region is more than 50% and 76%, respectively, smaller in the system with retroactivity compared to the one without it. Circuits, or portions of them, are often built on medium or high copy number plasmids, and even when built in a single copy, TFs still bind non-specifically to a large number of decoy sites [37, 38]. Therefore, unless retroactivity is mitigated, appropriately tuning the parameters is harder in practice than in an ideal modular system where the functionality of TFs is not affected by the downstream sites that they regulate. Also, this difficulty becomes more prominent as the circuit size increases. This is illustrated by the reduced robustness of the molecular signature classifier as compared to the event detector, and, in turn, by the reduced robustness of the event detector as compared to the toggle switch (Fig. 4, Fig. 8, Fig. 9, Fig. 12, Fig. 13).

As the circuit size increases, it is thus important to investigate ways of mitigating the effects of retroactivity. One avenue is the creation of insulation devices, which can be placed at suitable locations in the circuit to enable some level of modularity [22, 25]. Recent works have developed insulation devices for genetic circuits based on fast phosphorylation processes [25]. In fact, it was previously demonstrated theoretically that fast phosphorylation processes can be used to speed up the effective time scale of TFs, such that load-induced delays, occurring at the faster time scale of phosphorylation, become negligible on the slower time scale of gene expression [22]. This is consistent with our simulations showing that when the decay rate of TFs is artificially increased, proper behavior can be restored (Fig. 3, Fig. 6, Fig. 11).

Our focus here is the robustness of stability of equilibria as opposed to robustness of instability, such as found in oscillators [41]. In fact, in this case, the effects of retroactivity are not determined and have been shown to either increase or decrease the robustness of the oscillator design depending on the circuit topology [39].

We have provided an analytical approach to compare the robustness of stability of a system with retroactivity to that of a system without retroactivity using the concept of stability radius. This provides the largest perturbation a linear system’s matrix can tolerate before the appearance of eigenvalues with positive real part [29]. Since the stability radius is a tool developed for linear systems, we linearized the system about the steady state of interest and computed upper and lower bounds to the stability radius for both the system with retroactivity and the one without it. Comparisons among these analytical bounds lead to the finding that the system with retroactivity tends to have a smaller stability radius than that of the system without retroactivity, and hence a decreased robustness, confirming the numerical results.

In conclusion, our findings demonstrate that modularity leads to more robust systems in addition to having both evolutionary advantages in nature [21] and design advantages when engineering novel systems [40]. A modular approach to design, wherein the subsystems do not depend on their context, is therefore highly preferable to designs where large systems are monolithically created.

## Appendix

### Robustness Index: Stability Radius

Let Λ(*M*) denote the spectrum of a square matrix *M* ∈ 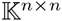, where 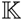 = ℂ or ℝ. Let ℂ^−^ denote the open left-half complex plane and let ℂ^+^ denote the closed right-half complex plane. Define the *stability radius* of *M* as

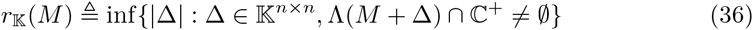

where | · | denotes the 2-norm. Then 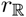(*M*) is the 2-norm of the smallest perturbation forcing *M* + Δ to be unstable. The stability radius defined in (36) is a natural measure of a system’s ability to maintain stability of an equilibrium point under perturbations to elements of the system matrix. A system with larger stability radius is able to maintain its stability under larger perturbations to the system’s matrix in the 2-norm sense. If Λ(*M*) ∩ ℂ^+^ ≠ 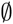, one has 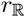(*M*) = 0. In the following, we only consider the non-trivial case: Λ(*M*) ∩ ℂ^+^ = 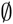, that is, *M* is a Hurwtiz stable matrix.

By the continuity of eigenvalues of a matrix with respect to its entries, the eigenvalue leaving ℂ^−^ towards ℂ^+^ must lie on ∂ℂ^−^, which is the boundary of ℂ^−^. Thus we can write

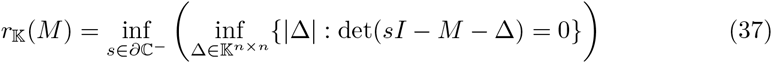

According to [29], one has the following relationship for the *complex stability radius* for Δ ∈ ℂ^*n × n*^:

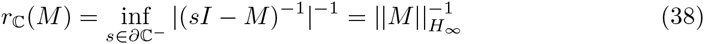

where

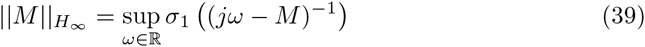

with *σ*_1_(·) the largest singular value of a matrix. This makes the computation of *r*_ℂ_(*M*) possible. By [30] one has that the real stability radius for Δ ∈ ℝ^*n × n*^ is given by

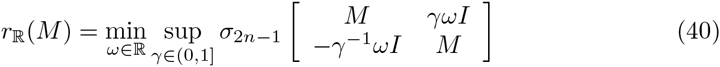

The real stability radius can be computed from (40), or by algorithms proposed in [31]. However, the computation of *r*_ℝ_(*M*) involves the minimization of unimodal functions [30], which is a challenging problem [32]. To avoid complex computations, we determine lower and upper bounds of *r*_ℝ_(*M*) which can be easily computed.

**Lemma 1** *Suppose M*, Δ ∈ ℝ^*n × n*^. Then

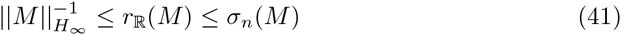

*where σ_n_(M) denotes the smallest singular value of M*.

**Proof of Lemma 1:** By the definition of the stability radius in (36), one immediately has *r*_ℂ_(*M*) ≤ *r*_ℝ_(*M*), which together with (38) implies the following lower bound

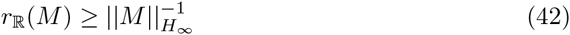

On the other hand, (37) implies

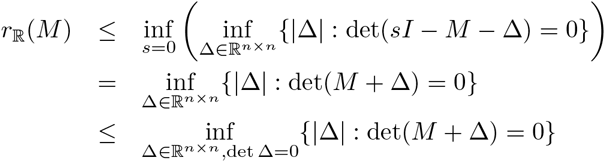

which is equal to *σ_n_(M)* by the Schmidt-Mirsky Theorem [33]. Then one has the following upper bound

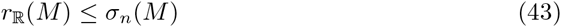

We complete the proof.

The bounds obtained in Lemma 1 are tight in the sense that they can be reached under certain conditions as indicated by the following lemma:

**Lemma 2** *If the H_∞_ norm of M is achieved at ω* = 0, *one has*

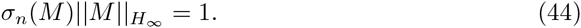

**Proof of Lemma 2:** Since the *H*_∞_ norm of *M* is achieved at *ω*,

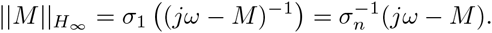

Then

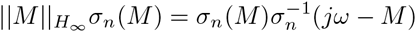

which is equal to 1 at *ω* = 0. We complete the proof.

Note that Lemma 1 has given upper and lower bounds for the real stability radius of a matrix. Such bounds have been shown to be tight in Lemma 2 in the sense that they can be reached under certain conditions. This in turn provides a simpler way to compare the robustness of linear systems as will be seen later in the following section.

### Robustness Comparison

For genetic networks in practice, parameter perturbations are usually real. If the real stability radius of 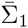 is larger than that of 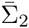, we say Σ_1_ is *more robust* than Σ_2_ at their equilibrium. By definition of real stability radius, this means that for all real parameter perturbations with a certain upper bound in its 2-norm, the system 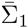 is stable at *x*\* while 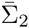 may become unstable. Note that the dynamics of Σ_i_ is similar to its linearized system 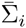 when it is sufficiently close to its equilibrium. Thus we call a non-linear system Σ_*i*_ is *more robust* if its linearized system 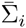 has larger real stability radius. In this subsection we will compare the robustness of the two linearized systems 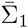 without retroactivity and 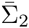 with retroactivity under real perturbations to elements of their system matrices *A* and (*I* + *R*)^−1^ *A*, respectively, by comparing their real stability radiuses. Since the real stability radius of an unstable matrix is 0, we only consider the case when *A* and (*I* + *R*)^−1^ *A* are both Hurwitz stable.

Based on Lemma 1, one can immediately conclude that Σ_1_ is more robust than Σ_2_ if

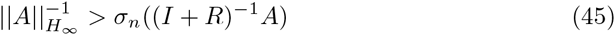

and Σ_2_ is more robust than Σ_1_ if

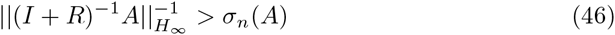

These two inequalities (45) and (46) give sufficient conditions to determine whether retroactivity increases or decreases the robustness of the gene transcription network against parameter perturbations.

When it comes to independent bindings, the retroactivity matrix *R* is diagonal [18], which allows us to obtain further analytical results. Let 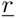 and 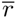 denote the smallest and largest diagonal entry of *R*. Then

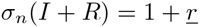

We further suppose that each node has at least one child. Then 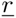 > 0 and thus

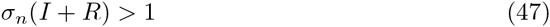

**Lemma 3** If

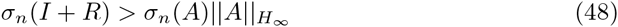

*one has that* Σ_1_ *is more robust than* Σ_2_.

**Proof of Lemma 3:** Let 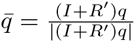, where *q* is the unit vector such that q′AA′*q* = λ_min_(AA′). Then

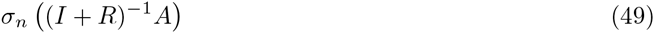

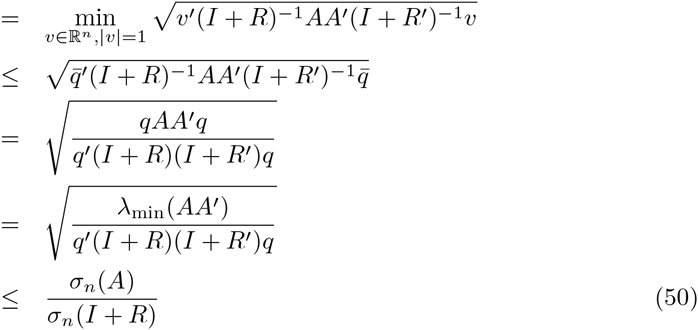

which and (48) imply

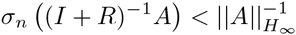

Then by Lemma 1 one has *r*_ℝ_((*I* + *R*)^−1^ *A*) < *r*_ℝ_(*A*).

It is worth mentioning that the condition in Lemma 3 separates the retroactivity matrix *R* and the system matrix *A*. From (47) and (50) one has

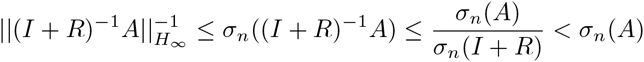

which implies that the condition (46) can not be satisfied in the case of independent bindings. Numerical computations suggest that (45) holds in general, the proof of which is quite challenging though. In the following we will look at serveral cases:

**Case 1:** Assume that retroactivities corresponding to all TF/promoter bindings in a gene transcription network are balanced in the sense that 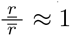 1. Let

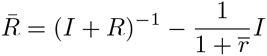

By (37), one has

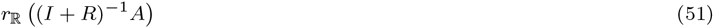

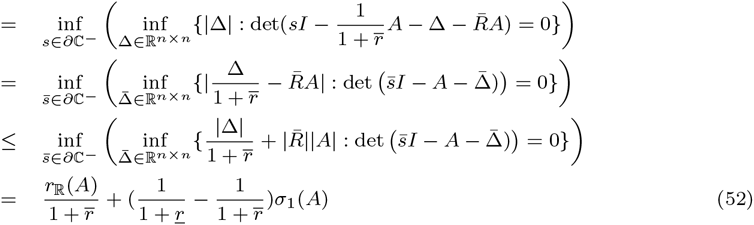

Since 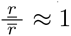, then

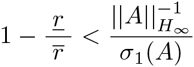

It follows that

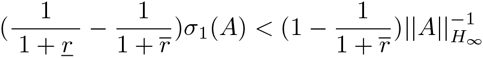

from which, 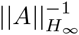 ≤ *r*_ℝ_(*A*) and (52), one has

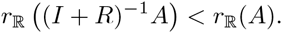

Then Σ_1_ is more robust than Σ_2_.

**Case 2:** Assume that there exists one TF/promoter binding which leads to extremely large retroactivity, or in other words, 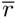 → ∞. Note that

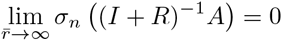

By the continuity of eigenvalues of a matrix with respect to its entries, there must exist a finite real number μ such that for all 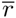 ∈ [μ, ∞), *σ_n_* ((*I* + *R*)^−1^ *A*) < 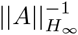. Then Σ_1_ is more robust than Σ_2_.

**Case 3:** Assume that the retroactivity corresponding to each TF/promoter binding in a gene transcription network is sufficiently large. Since *A* is Hurwtitz stable, one has ∥*A*∥_*H*_∞__ and *σ_n_(A)* are bounded. Then when 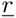 is large enough, one has (48).

**Case 4:** Assume that the *H*_∞_ norm of the matrix *A* is achieved at *ω* = 0, which in practice suggests that Σ_1_ has a “low-pass filter” behavior. Because of its benefit to ignore rapid variations and only respond to longer-lasting changes, this low-pass filtering capacity is a common feature of regulation of transcription, as suggested in *E*. *coli* theoretically [34], verified experimentally [35] and recently observed in eukaryotes [36]. By Lemma 2 on has ∥*A*∥_*H*_∞__ *σ_n_(A)* = 1. Note that *π_n_*(*I* + *R*) = 1 + 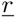 > 1. Then (48) holds, which implies Σ_1_ is more robust than Σ_2_.

As a summary of the above findings, we have the following theorem

**Theorem 1** In the case of independent bindings, let 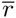 and 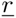 denote the largest and the smallest diagonal entry of R, respectively. Σ_1_ is more robust than Σ_2_ if any of the followings holds:

- retroactivities corresponding to all TF/promoter bindings in a gene transcription network are balanced in the sense that 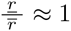;
- there exists one TF/promoter binding which leads to extremely large retroactivity in the sense that 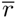 ≈ ∞;
- the retroactivity corresponding to each TF/promoter binding in a gene transcription network is sufficiently large in the sense that 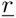 is large;
- the *H*_∞_ norm of the matrix A is achieved at ω = 0.

## Supporting Information

### S1 Fig

Supplementary figures to show the percentage numbers appearing in the pie figures are the values that simulations results converge to.

